# VideoTagger: User-Friendly Software for Annotating Video Experiments of Any Duration

**DOI:** 10.1101/272468

**Authors:** Peter Rennert, Oisin Mac Aodha, Matthew Piper, Gabriel Brostow

**Author notes:** To whom correspondence should be addressed: MP - GB.

## Abstract

**Background:** Scientific insight is often sought by recording and analyzing large quantities of video. While easy access to cameras has increased the quantity of collected videos, the rate at which they can be analyzed remains a major limitation. Often, bench scientists struggle with the most basic problem that there is currently no user-friendly, flexible, and open source software tool with which to watch and annotate these videos.

**Results:** We have created the VideoTagger tool to address these and many of the other associated challenges of video analysis. VideoTagger allows non-programming users to efficiently explore, annotate, and visualize large quantities of video data, within their existing experimental protocols. Further, it is built to accept programmed plugins written in Python, to enable seamless integration with other sophisticated computer-aided analyses.

We tested VideoTagger ourselves, and have a growing base of users in other scientific disciplines. Capitalising on the unique features of VideoTagger to play back infinite lengths of video footage at various speeds, we annotated 39h of a *Drosophila melanogaster* lifespan video, at approximately 10-15x faster than real-time. We then used these labels to train a machine-learning plugin, which we used to annotate an additional 538h of footage automatically. In this way, we found that flies fall over spontaneously with increasing frequency as they age, and also spend longer durations struggling to right themselves. Ageing in flies is typically defined by length of life. We propose that this new mobility measure of ageing could help the discovery of mechanisms in biogerontology, refining our definition of what healthy ageing means in this extremely small, but widely used, invertebrate.

**Conclusions:** We show how VideoTagger is sufficiently flexible for studying lengthy and/or numerous video experiments, thus directly improving scientists’ productivity across varied domains.

## Background

Increasingly, people rely on video footage to support their scientific claims. Just recording the videos is ever easier, as cameras and memory get cheaper, smaller, and less power-hungry. However, this has created new problems: the sheer quantity of video data and their intricate formats frustrate otherwise hands-on researchers. Once the video material is recorded, we find that researchers typically face three main tasks: exploration, annotation, and visualization, that are hampered by the lack of easy-to-use software systems. There are several existing software tools to do annotation and classification, but their use is either restricted to users with programming experience and/or they are built to perform a narrow set of functions that restrict their use to some specific tasks [1],[2],[3]. We have also found that they are often not designed to handle large video datasets, which is problematic for many uses, such as behavior analysis and biodiversity surveys in ecology [4, 5]. Rather than repeating the cycle of developing a ground-up software solution for our specific task, we followed the guidelines in [6] to develop VideoTagger: an easy-to-install cross platform and modular open source tool, that provides an end-to-end solution to facilitate scientific analysis of video data. Similar to the image-focused software system ImageJ [7], VideoTagger stands alone or supports plugins that can be used for automated video processing or machine learning and statistical analysis. The software is easy to use and thus immediately accessible for biologists to undertake manual annotation tasks, while also providing a platform for more sophisticated, automated analyses with the use of machine learning routines. We illustrate the VideoTagger workflow with different use cases in the supplementary material (Supplementary Manual). Here, we demonstrate the system’s features using footage from an underwater habitat study, where a researcher was faced with the challenge of counting the number of fish species within a 2 hour video. We then demonstrate how VideoTagger can enable biological discovery in extremely long videos (>1000h continuous footage) by facilitating a cycle of training and automated action detection with a machine learning plugin.

## Implementation

Our tool, VideoTagger (available on https://github.com/groakat/pyTools), addresses the problems of exploring, annotating, and visualizing the annotations in videos of any length. VideoTagger was developed using Python and the Qt wrapper PySide. The implementation is cross platform (tested on Windows 10, OSX, and Ubuntu) and is designed with non-programming users in mind, as well as being expandable via plugins written in Python. Using the video library ffmpeg, we can decode virtually any video. Our data structures and algorithms make it possible to browse and annotate videos of any length and display the annotations in an interactive and intuitive manner. By working on a low-resolution version of the video, we achieve high browsing speeds (up to 60x), while allowing the user to seamlessly obtain a high-resolution frame when more detail is required. Multi-threaded caching allows VideoTagger to smoothly move forward and backward through video footage. Finally, any annotations made in VideoTagger can be exported to a .CSV file for further analysis. The following sections describe core-functionality in detail.

Further instructions on installation, running, and using the software can be found online (http://peter.rennert.io/papers/videotagger/install/) and in the supplementary material (Supplementary_material_02). We have also made an overview video to showcase the available functionality, as well as provided self-contained example files of use-cases featuring mice, flies, and fish, based on data from existing users.

## Results and Discussion

### Exploration: variable speed playback of a wide range of video formats of any duration

At any point in video data collection, users want to explore the recorded material to assess the occurrences of expected phenomena and/or survey for new ones. VideoTagger makes this possible for most standard video compression formats by building on the open source ffmpeg library for decoding and encoding video (Figure 1a).

**Figure 1:**
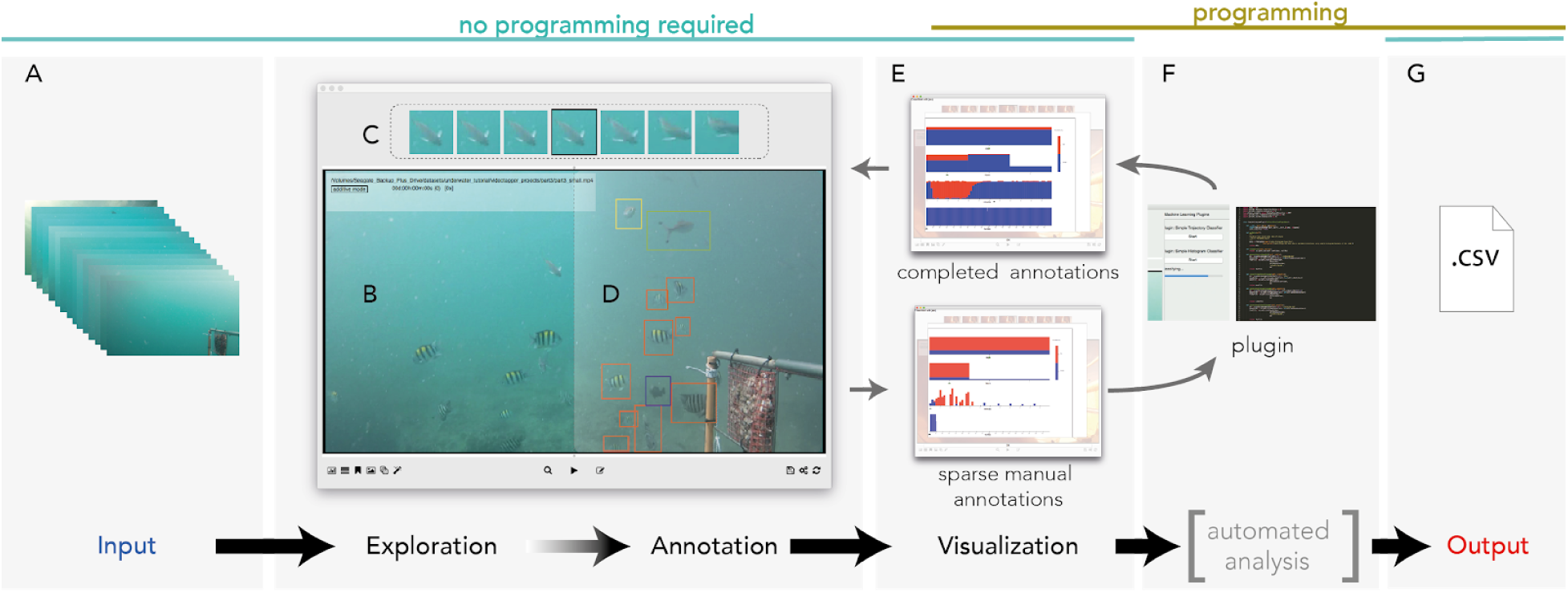
VideoTagger overview and workflow. Users can import a wide variety of video formats (A), which are displayed on the main viewing frame (B & D). In the example above, the screen is split to represent the initial exploration phase (B) from the annotation phase (D). Above the main view is a panel of thumbnails showing the previous three frames, the current frame and three future frames (C) that facilitate observation of the beginning and end of a focal event. Action buttons for annotations, view options, search, save and settings adjustments can be found in the lower panel. To visualize annotated events, an interactive multi time-scale hierarchical view is generated in a separate window (E). An option also exists to integrate user-developed plugins (F), for example to enable automatic detection of actions in the footage. All annotations can be exported to an easy to read format for use in further analyses (G). Importantly, step (F) is optional such that a complete workflow can be undertaken without programming expertise.

While surveying video data, speed is important, because real-time playback (at 1x) is too slow for some segments, and too fast for others. In the case of the underwater habitat study, a remote camera was set to record footage of fish attracted by bait. We expected to see a constant stream of many fish, however, during the exploration phase (Figure 1b) it became apparent that long sections of the videos were empty sea, and only brief periods featured intense feeding activity by many fish. VideoTagger facilitates exploration by enabling playback speeds to range from single frame steps up to 60 times the captured frame rate of 25fps. It does this by generating and displaying a reduced-resolution version (640px across by an appropriate resolution to preserve aspect ratio) of each video. This remains linked to the original footage, so that the high resolution frame can be retrieved at any time. In addition, the interface displays an array of pre-view and post-view thumbnails (Figure 1c), to make it easier to see the start or end of an approaching event when viewing the video at speed. To modulate playback speed, the interface works with a keyboard and mouse, but also supports adding an inexpensive USB jog-shuttle device for fast video navigation.

For efficient exploration of long video sequences, VideoTagger chains smaller videos into a seamless larger one by pre-loading subsequent files, with no limit on the duration of concatenated videos. Consequently, footage spanning days and months can be captured and stored as small video chunks, making storage and backup more robust, and simplifying distribution of raw data.

### Annotation: labelling tools to manually highlight events of interest

After gaining a qualitative impression of the data, users may wish to note specific events of interest (Figure 1d). We have therefore provided four different labelling tools that, in our experience, cover the requirements of most users who wish to annotate events in long videos. In our underwater example, users can 1) bookmark a moment in the timeline for sharing or revisiting later, 2) note the presence or absence of fish generally, on a whole-frame basis, 3) annotate each individual fish in a frame with a separate bounding box and associated label, with no maximum limit on annotations, and/or 4) follow the location and size of a fish as it moves over time by using the mouse and its scroll-wheel to adjust the position and size of its associated bounding box. Thus, the user can easily follow an object of interest through the footage, without diverting their attention from it, to retreat or advance along the timeline.Annotations are tagged with the person’s name, allowing for easy consistency-checks and collaboration.

### Visualization: an interactive hierarchical view of event labels

A visual summary of annotations is important to gain an overview of notable events in the footage. However, tagged events can potentially span the whole length of a video, ranging from seconds to months in length. This creates a scale problem in the visualization, since fleeting observations, like shark sightings in our ocean footage, would be imperceptibly small and therefore lost in a simple linear timeline representation. We address this problem by displaying an interactive hierarchical view, which combines a zooming-paradigm with time-series information visualization practices [8] (Figure 1e). The top row of the hierarchy conveys the presence or absence of annotated events at the time scale of whole days, in the form of a histogram using near logarithmic proportions. Clicking on a particular day selects it and changes the next level of visualized detail on the row beneath to show an hour-by-hour display of that day’s annotated events. In the same way, subsequent rows respectively show per-minute and per-frame annotations. Clicking on a frame triggers the main video display to jump to that moment in the overall video. This allows for quick inspection of complex annotations in long videos, at whatever time scale is convenient.

We illustrate the hierarchical view for our underwater example (Figure 1e): we annotated a portion of the frames in the footage as containing fish or no fish. As the visualization is interactive, these labels can be surveyed by another user, possibly with greater expertise, who can click and jump directly to time-points deemed to contain fish, to identify the distinct species. By enabling such collaboration, a species-level ecological survey can be achieved with much greater efficiency.

### Plugins: interface to expand input/output and machine learning functionality

The amount of video material a researcher can view and annotate by hand is limited by the time available. In many cases, the task is very repetitive. There, an automated machine learning solution could relieve part of the manual burden. Similar to ImageJ [9], VideoTagger provides a plugin interface which exposes the video content and annotations to third-party software systems and user created Python code. We are working toward establishing a library of machine learning plugins to provide standard off-the-shelf solutions, and encourage practitioners that use VideoTagger to make their plugins available for other researchers in our public code repository. In the underwater example, once the presence of fish has been annotated in one video, a classifier is trained and applied to the remaining 200 videos of the study to quickly quantify the total time fish are present (Figure 1e). By populating the interactive hierarchical view with these automated annotations, species identification is further accelerated on a larger scale by enabling the expert to skip over large amounts of uninformative footage.

VideoTagger can read in annotations from other software (e.g. CTRAX [2]). Further, all annotations, settings, and bookmarks are stored in CSV files that can easily be imported into other programming and scripting packages. This enables pre- and post-processing of the annotation data outside VideoTagger, and integration into existing workflows.

### **Biological Ageing in *Drosophila***

Biogerontology seeks to uncover genetic and environmental manipulations that enhance healthy ageing. Since demonstrations that the mechanistic basis of ageing is evolutionarily conserved [10], many important discoveries have been made through experiments on short lived, genetically well characterised model organisms such as the fruitfly *Drosophila melanogaster*[11]. Female flies are ∼2.5mm long, which has enormous cost advantages for laboratory rearing of large populations, but comes with the downside that observing physical and behavioral health is far from simple. This raises the concern that while we may discover interventions that increase lifespan, they may not enhance healthspan, thus inadvertently promoting an extension of the time spent moribund. We took advantage of VideoTagger’s ability to access extremely long-term footage to survey ∼78 days of near-continuous video of flies as they aged. In an initial survey, we noticed that aged flies were more prone to falling onto their backs spontaneously and spending time struggling to get up (Figure 2a). This appeared to be similar to the supine behavior reported for old Medflies [12] and which had been reported as absent in relatively sparse observations of *Drosophila* lifespans [13]. We annotated footage for struggling using the eleventh hour of each of the first 42 days of several flies’ lifespans. In total, these 42 hours of footage took ∼4 hours to annotate, and we found that the duration of time the flies spent struggling increased dramatically as they aged, presumably due to worsening neuronal and/or motor functioning (Figure 2b).

**Figure 2:**
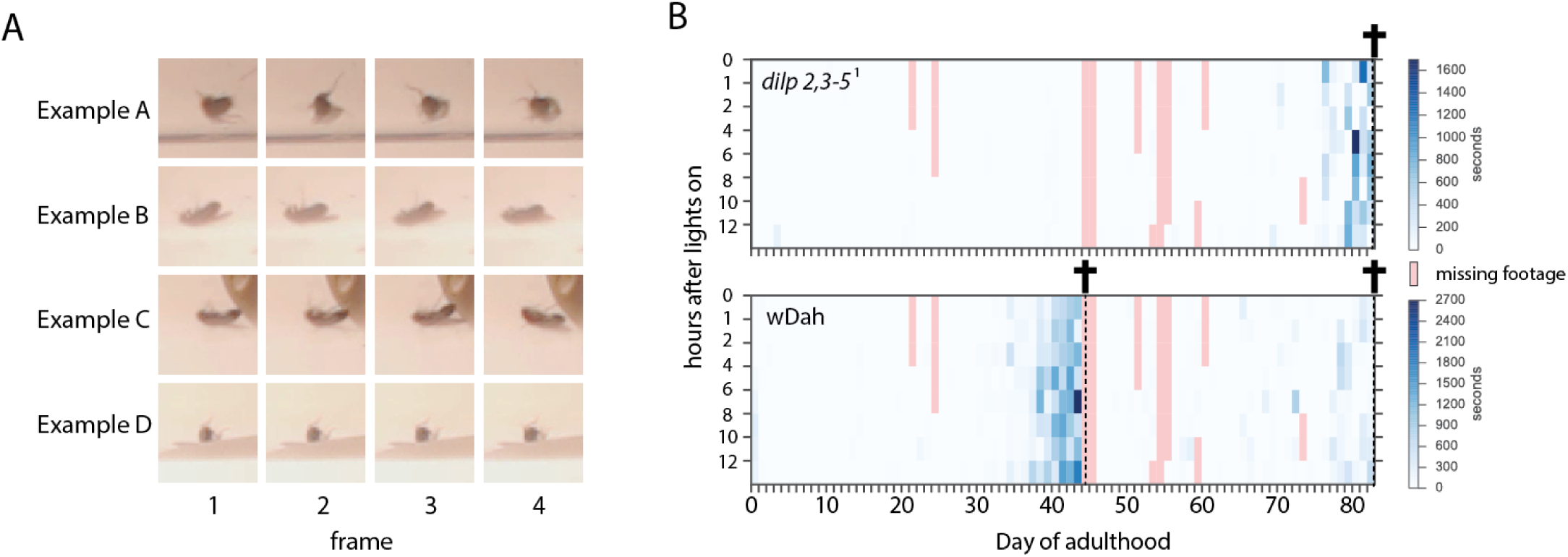
Struggling in ageing flies. (A) Example footage of flies struggling to right themselves. Although we only show four frames per sequence, in practice the duration of an individual struggling event can range from milliseconds to minutes. (B) We manually scored struggling for one hour of each day of life during the lifespans of long-lived mutant (*dilp2,3-5*^1^) and wild type (wDah) flies. Additional instances were scored using a machine learning plugin that achieved 93% accuracy. As flies approached death, the time spent struggling increased, for all times throughout the day. This late-life period of altered physical function was not increased in the mutant, relative to controls, indicating they benefitted from both longer life and healthspan. Data for three flies (one mutant and two controls) shown. Day of death is indicated by crosses and intensity of blue in each 1h block indicates amount of time spent struggling. Note that the range of blue scales for the two panels differs such that the wild type flies had more instances of struggling than the mutant.

To extend these annotations, we wrote a plugin that learned from manual annotations and automatically labeled further instances. Our plugin automatically detects the position of a fly in a vial, links those detections to create continuous tracks, and finally determines if a fly is struggling using both appearance and motion features. More details of this plugin can be found in our supplementary material. For evaluation, we trained the behavior classifier on even numbered days of the 42 day manual annotations and assessed its accuracy by comparing automated predictions with manual annotations for odd days. Given the plugin’s high degree of accuracy (93%; Figure S3), we extended the automated annotations to cover additional footage. Within the computing time we devoted to this project, we automatically predicted behavior for a total of eight hours per day for 78 days worth of footage (an additional 582 hours) for both long-lived insulin mutant flies (*w*Dah; *dilp2,3-5*^*1*^ null) and their wild-type controls (*w*Dah) [14].This revealed an increase in struggling instances for all times of day during the last 7-10 days of all the flies’ lives. Importantly, this worsening state was not extended in the long-lived mutant when compared with controls, despite their increased lifespan. Thus, similar to long-lived insulin mutant worms [15], the enhanced lifespan of insulin mutant flies appears to be accompanied by a longer period of better physical health.

This example serves to illustrate how VideoTagger can expose thousands of hours of continuous footage to a combination of human and automated analysis for detection of relatively rare actions that can be biologically informative. We propose that spontaneous falling and struggling, a behavior that we observed for only 2.5% of the lifetime of our wild type (wDah) flies, may be a simple and good indicator of age-related health that, with automated detection, could be used to assess if some of the well established genetically long lived fly models also retain youthful mobility for longer. Further, since falling and struggling are unprovoked actions, it could be a low effort marker to screen for new genes that confer prolonged health, whether or not it is accompanied by longer lifespan.

## Conclusion

We have described a software tool that meets the general requirements of laboratory scientists to make a quick start on surveying and annotating their video data. It has a simple appearance, is cross-platform, is easy to install and use (see supplementary material Manual), but is also sufficiently feature-rich and powerful to facilitate more complex computer-aided video analysis tasks. By providing the platform from which users can build domain specific solutions, we solve the initial problem that a significant overhead is required to make state of the art machine learning algorithms accessible for end users who rely on video data. We propose that by removing this barrier, VideoTagger facilitates biological discovery and facilitates collaborations between biologists and computer scientists.

## Methods

### ***Drosophila* handling and rearing**

We used insulin ligand mutants *w*Dah *dilp*2,3-5 and their wild-type controls *w*Dah as described in [14]. All flies were reared at standard density on 1x sugar / yeast medium (1SY) as described in [16]. Throughout their lifespans, flies were transferred to fresh food 3 times a week, at which point deaths were scored. Individual flies were removed from the population reared for lifespan and maintained separately for recording. Lifespans were maintained at 25C, 65% humidity, 12h:12h light:dark cycle.

### Recording housing and equipment

Fish videos

The videos used for the fish classification experiment were recorded with a GoPro camera in a waterproof housing. Together with a bait, the camera was attached to a buoy and submerged in the ocean. The setup filmed passively for about 2h after which the camera was retrieved and the data copied.

#### Fly videos

To record the *Drosophila* life-spans, we built a custom light-proof housing which could fit 4 standard vials. The housing was fitted with custom made diffused background illumination which can be switched remotely by software from night mode (infrared LEDs) to day mode (white LEDs). As a recording device we used a Logitech C920 webcam with the IR filter removed, from which we grabbed the H264 stream and saved it in one-minute intervals.

### VideoTagger development

The design of VideoTagger was focused on allowing precise and fast access to video frames and giving the user various tools to annotate and review annotations of sections of videos.VideoTagger was developed using Python and the Qt wrapper PySide. This choice allowed us to develop a cross-platform native application which can be easily extended by adding Python modules to it. Using Python also makes it easy to reuse parts of VideoTagger (e.g. the video interface) for other scientific projects, for instance to write sophisticated plugins. Distribution of files for installation on different platforms and user guides can be found in the Supplementary Material and at http://peter.rennert.io/papers/videotagger/install/.

#### Abbreviations

CSV: comma separated files
wDah: white Dahomey
Dilp: *Drosophila* insulin-like peptide
SY: Sugar / Yeast (food)
IR: infra-red
PCA: principal component analysis

## Declarations

**Ethics approval and consent to participate**

Not applicable

## Consent for publication

Permission has been sought and granted for footage used to demonstrate VideoTagger: mouse data was sourced from [4, 17] and fish/ocean footage was accessible via the Centre for Marine Futures at the University of Western Australia [18]

## Availability of data and material

The source code is Open Source on GitHub. The 3-month long video footage of flies that was generated and analysed in this study is available from the corresponding author on request. Adequate storage will need to be provided (∼20 TeraBytes of hard disks).

## Authors’ contributions

Conception: all authors; Data collection: PR; Data analysis: all authors; Wrote the manuscript: all authors

## Acknowledgements

The authors would like to thank David Curnick and Gioia De Franceschi for testing the alpha version of VideoTagger. We also thank the Centre for Marine Futures at the University of Western Australia for access to the fish footage.

**Authors’ information (optional)**

